# Altered development and network connectivity in a human neuronal model of 15q11.2 deletion-related neurodevelopmental disorders

**DOI:** 10.1101/2024.09.19.613912

**Authors:** Christa W. Habela, Shiyu Liu, Arens Taga, Raha Dastgheyb, Norman Haughey, Dwight Bergles, Hongjun Song, Guo-Li Ming, Nicholas J. Maragakis

## Abstract

The chromosome 15q11.2 locus is deleted in 1.5% of patients with genetic epilepsy and confers a risk for intellectual disability and schizophrenia. Individuals with this deletion demonstrate increased cortical thickness, decreased cortical surface area and white matter abnormalities. Human induced pluripotent stem cell (iPSC)-derived neural progenitor cells (NPC) from 15q11.2 deletion individuals exhibit early adhesion junction and migration abnormalities, but later neuronal development and function have not been fully assessed. Imaging studies indicating altered structure and network connectivity in the anterior brain regions and the cingulum suggest that in addition to alterations in progenitor dynamics, there may also be structural and functional changes within discrete networks of mature neurons. To explore this, we generated human forebrain cortical neurons from iPSCs derived from individuals with or without 15q11.2 deletion and used longitudinal imaging and multielectrode array analysis to evaluate neuronal development over time. 15q11.2 deleted neurons exhibited fewer connections and an increase in inhibitory neurons. Individual neurons had decreased neurite complexity and overall decreased neurite length. These structural changes were associated with a reduction in multiunit action potential generation, bursting and synchronization. The 15q11.2 deleted neurons also demonstrated specific functional deficits in glutamate and GABA mediated network activity and synchronization with a delay in the maturation of the inhibitory response to GABA. These data indicate that deletion of the 15q11.2 region is sufficient to impair the structural and functional maturation of cortical neuron networks which likely underlies the pathologic changes in humans with the 15q11.2 deletion.

## Introduction

Neurodevelopmental and neuropsychiatric disorders such as epilepsy, schizophrenia, autism and intellectual disability can be acquired but are most commonly caused by genetic changes that may be inherited or occur *de novo*. Among inherited forms copy number variations (CNV’s) are common^1–4^. These can involve large deletions or duplications affecting many genes and regulatory sequences making mechanistic study and predictions difficult using standard models. However, the use of induced pluripotent stem cell derived cells (iPSCs) from individuals bearing these changes provides a mechanism to uncover changes responsible for altered neuronal development and excitability. A CNV in the 15q11.2 region in humans is implicated in both neurodevelopmental and neuropsychiatric disorders^5–9^. Microdeletion between breakpoints 1 and 2 results in haploinsufficiency of 4 non-imprinted genes adjacent to the imprinted regions involved in Prader Willi and Angelman syndrome, nonimprinted in Prader-Willi/Angelman syndrome-1 (*NIPA1*), nonimprinted in Prader-Willi/Angelman syndrome-2 (NIPA2), tubulin gamma complex-associated protein-5 (*TUBGCP5*) and Cytoplasmic fragile X mental retardation-interacting protein-1 (*CYFIP1*)^10^. This microdeletion is found in up to 1.4% of patients with generalized epilepsy, making it one of the most common CNVs identified in epilepsy^3, 8, 11^. 15q11.2 haploinsufficiency also confers risk for schizophrenia and intellectual disability^1, 5, 6, 12^. Confounding the characterization of the effects of the deletion on development is the fact that in families which have multiple family members with the deletion, there is significant heterogeneity in the presentation, with some severely affected children inheriting the deletion from more mildly affected or unaffected parents^5^. However, independent of neuropsychiatric diagnoses, the 15q11.2 microdeletion has been shown to affect overall IQ^6^ as well as the presence of dyscalculia and dyslexia^13^.

Structural analyses of brain magnetic resonance imaging studies (MRIs) of individuals with the 15q11.2 microdeletion have revealed reduced surface area and increased cortical thickness predominately in the frontal lobe, anterior cingulate and the precentral and postcentral gyri, as well as decreased volume of the nucleus accumbens compared to controls, even in the absence of clinically apparent disease^14^. Analysis of white matter structure using diffusion tensor imaging (DTI) studies demonstrates that 15q11.2 deletion carriers also have increased fractional anisotropy (FA) in the internal capsule and the hippocampal portion of the cingulum and decreased FA in the posterior thalamic radiation compared to controls, with the latter being associated with worse cognitive function^15^. Alterations in white matter tract microstructure detected by DTI can reflect changes in myelination or in the structure, packing density or branching of axons^16^. It is unknown which process is represented by the DTI changes in individuals with 15q11.2 deletions and the developmental and cellular mechanisms that lead to them are unknown.

Human induced pluripotent stem cells (hiPSCs) harboring the 15q11.2 microdeletion exhibit dysregulation of adherens junctions in neural progenitor cells (NPCs) in early neuronal development^17^. This has been attributed specifically to haploinsufficiency of *CYFIP1,* one of the four genes on this locus. Notably, deletion of *Cyfip1* in mice results in ectopic embryonic NPCs^17^, as well as disruption of the adult neurogenic niche in the subventricular zone and altered proliferation of NPCs in adult mice^18^. CYFIP1 interacts with the WAVE regulatory complex that functions in actin nucleation and polymerization required for cell motility, cell polarity, neurite outgrowth and synaptic remodeling ^19–24^. It also binds the Fragile X mental retardation protein (FMRP) and is required for FMRP-mediated activity-dependent translation repression at synapses^25, 26^. Loss of *Cyfip1* in rodents leads to multiple behavioral changes that are consistent with the positive and negative symptoms of schizophrenia^27–30^. Further, loss of *Cyfip1* leads to alterations in white matter structure in rodents^29, 30^. This phenotypic overlap between CYFIP1-deficient rodents and 15q11.2 deletion harboring humans, as well as impaired neurogenesis in 15q11.2 deletion iPSCs, suggests that loss of *CYFIP1* may be a primary contributor to the alterations in neuronal activity in these patients. However, the structure and function of 15q11.2 deletion neurons in later stages of development have not been determined.

Given the variable phenotypes and incomplete penetrance in human individuals with 15q11.2 deletions, a fundamental question is whether this deletion alone is sufficient to bring about the altered neuronal activity and connectivity present in patients with intellectual disability, schizophrenia and epilepsy. To address this question, we used confocal microscopy and multielectrode array (MEA) analysis to examine the structural and functional development of human iPSCs derived from three 15q11.2 deletion harboring donors as well as six healthy controls without 15q11.2 deletion over extended periods of time in culture. The results indicate that 15q11.2 deletion neurons have structural changes including decreased neurite arborization, an increase in the proportion of inhibitory neurons within cultures and deficits in electrophysiologic maturation. We hypothesize that such changes predispose 15q11.2 deletion individuals to the macroscopic structural changes and the neuropsychiatric phenotypes of 15q11.2 deletion individuals

## Methods

### Differentiating Cortical NPCs from hiPSCs

Human induced pluripotent stem cells (hiPSCs) were maintained in Essential 8 media (ThermoFisher Scientific, Waltham, MA, USA) as described previously^31^. hiPSC derived cortical neural progenitor cells (NPCs) were generated from either control or 15q11.2 deletion bearing patient lines (**Supplementary Table 1**) using the human forebrain cortical differentiation protocol as adapted from Wen et al 2014^32^. On day in culture (DIC) 1, hiPSC plates were cleaned and were lifted from the plate with 1 mg/mL collagenase (ThermoFisher) in DMEM/F12 media (Corning, Corning, NY, USA) for 1 hour at 37°C. The colonies were transferred to a 15 mL conical tube, allowed to settle to the bottom of the tube by gravity and the media was aspirated. Colonies were then gently resuspended in DA media (DMEMF12 (-) L – Glutamine (Corning), 20% Knockout Serum Replacement, 1:100 Glutamax, 1:100 Pen-Strep, 1:100 Non Essential Amino Acids (NEAA), 1:1000 β-mercapthoethanol (ThermoFisher), 2µM Dorsomorphin, 2µM A83-01 (StemCell Technologies, Cambridge, MA, USA)) supplemented with 20 µM Rho-associated coiled-coil forming protein serine/threonine kinase inhibitor (ROCK-I, compound Y-27632) (ThermoFisher) and transferred to a 6-well ultra-low attachment plate (Corning) to form embryoid bodies (EBs) in suspension. On DIC 2- 4, EBs were washed and media was fully changed with DA media. On DIC 5, media was changed to NPC media (DMEMF12 (-) L - Glutamine, 1:100 Glutamax, 1:100 N2 supplement (ThermoFisher), 2 µM Cyclopamine (StemCell Technologies), 2 µg/mL Heparin) for 48 hours. On Day 7, the EBs were transferred in fresh NPC media to plates coated with 2% Matrigel (Corning) for 24 hours at 37°C. Partial NPC media exchanges were performed every other day until DIC 21. On DIC 22, the resulting neural rosettes were washed with phosphate buffered saline (PBS) and incubated in StemDiff dissociation media (ThermoFisher) for 1 hour at 37°C. Rosettes were lifted using pipet pressure application of pre-warmed DMEM-F12 directed at the center of the rosette and transferred to a six-well ultra-low adhesion plate and incubated overnight (16-20 hours) at 37°C in NPC media supplemented 1:100 with B27 Supplement (ThermoFisher). On DIC 23, rosettes were transferred to StemPro Accutase media (ThermoFisher) for 10 minutes at 37°C before trituration to dissociate to single cells followed by centrifugation at 350 x g for 5 minutes. NPCs were pelleted and resuspended in neuronal media (Neurobasal media (GIBCO-ThermoFisher Scientific), Glutamax, nonessential amino acids, B27 supplement, 10 ng / mL BDNF and 10 ng / mL GDNF (Preprotech – ThermoFisher) and transferred to cryovials containing DMSO (Millipore Sigma, Darmstadt, Germany) for a final DMSO concentration of 10% DMSO. Cryovials were transferred to -80 °C in sealed Styrofoam for a minimum of 24 hours prior to storage in liquid nitrogen.

### Neuronal differentiation

Liquid nitrogen stocks of DIC 23 NPCs were diluted in 10 mL prewarmed Neurobasal media after brief incubation in a 37°C water bath, centrifuged at 350 x g for 5 minutes and resuspended in 10 mLs of neuronal media supplemented 20 μM of ROCK-I (1:500 DMSO) and plated on a 10 cm plate coated with poly-L-ornithine (PLO) and laminin as previously described^31^. After 48 hours, ROCK-I was removed, and Neuronal media supplemented with 125 nM of the notch inhibitor, Compound E (StemCell Technologies Vancouver, Canada) was exchanged every 48 to 72 hours with every other media change containing 1 ug/mL laminin (ThermoFisher). After 7- 10 days differentiation, neuronal plates were treated for 48 hours with 0.2 uM cytosine arabinoside (ARA-C) in neuronal media to reduce actively proliferating cells.

### Neuron and astrocyte co-culture

DIC 37 to 45 neurons and cortical rat astrocytes at passage 7 to 8^33^ were washed once in PBS and then lifted with 0.05 % trypsin (ThermoFisher), centrifuged at 350 x g for 5 minutes and resuspended in 1 mL neuronal medium supplemented with compound E, 1:100 antibacterial – antifungal (Anti-Anti) (ThermoFisher) and 5% fetal bovine serum (FBS) (MEA-FBS media) with 20 µM ROCK-I. Neurons were counted and resuspended at a density of 10,000 cells / µL and astrocytes resuspended at 5,000 cells / µL. For multielectrode array experiments, neurons and astrocytes were combined at a volume ratio of 1:1 (cell count ratio of 2:1) and plated as a 5 µl droplet to the center of a PLO and laminin-coated electrode array in an Axion Lumos 24-well MEA plate (Axion Biosystems, Atlanta, GA, USA). For imaging experiments, cells were plated at 50,000 neurons and 25,000 astrocytes per coverslip onto washed 30 mm #1 German borosilicate coverslips coated with PLO and laminin per well of a 24-well plate. For each coverslip, 10 µL of the cell suspension was mixed with 40 µL of MEA media with 20 µM ROCK – I and placed on the center of the coverslip. After a 30-minute incubation at 37°C 5% CO_2_, the wells of both the MEA plates and the 24-well coverslip plates were flooded with 500 µL MEA- FBS media added in 2 – 250 µL parts to each side of the well.

### Cell Transfection

Plasmid DNA from the pAAV-hSyn-EGFP construct (Addgene, Watertaown, MA, USA) was diluted into 50 µL of Opti-MEM Reduced Serum Media (ThermoFisher) and mixed gently. Lipofectamine 2000 (ThermoFisher) was mixed into another 50 µL of Opti-MEM medium and incubated for 5 minutes at room temperature. The diluted DNA and diluted Lipofectamine 2000 were mixed gently and incubated at room temperature for 20 minutes. 100 µL of the mixture was added to each well and incubated in a 37°C incubator for 24 hours before a complete media change was performed. Neurons were transfected 3 to 7 days after co-culture.

### Multi-electrode array (MEA) recording and drug treatment

MEA recordings were obtained on a Maestro Edge multi-well MEA system (Axion Biosystems) with environmental control set to 37°C, 5% CO2. On the day of recording plates allowed to equilibrate for 5 minutes prior to recording. For longitudinal recordings of spontaneous activity, plates were recorded for 5 minutes. The entire 5-minute record was used for analysis. Raw voltage was sampled at 12.5 Hz and filtered with a 200 Hz high pass and 3kHz low pass Butterworth filter. Spikes were defined as adaptive threshold crossings of greater than six times the standard deviation. Bursts were defined by a minimum of 5 spikes on a single electrode with a maximum inter spike interval (ISI) of 100 ms and network bursts were defined by an envelope threshold with a threshold factor of 1.25 with a minimum inter-burst interval of 100 ms, minimum of 35% of electrodes included and 75% of bursts^34^. Weighted mean firing rate (WMFR) (the mean firing rate averaged across active electrodes), the burst percentage (the number of spikes occurring in bursts divided by the total number of spikes multiplied by 100) and synchronization (area under normalized cross correlation - AUNCC)^35^ were analyzed. For drug application, a minimum of 2 minutes baseline recording was obtained prior to manually pipetting 20 µL of either warmed media or warmed media with DMSO vehicle while continuous recording continued for 3 minutes prior to adding 20 µL of warmed media containing sufficient drug to achieve the final desired concentration of each drug. Recording continued for another 3 minutes prior to removing plates and washing out drugs. The drug-dependent effects were determined by the percent change in activity compared to the vehicle application. Bicuculline (Millipore-Sigma) was diluted in DMSO to a stock concentration of 20 mM and used at a final concentration of 10 µM (1:2000 DMSO dilution). CNQX (6-cyano-7- nitroquinoxaline-2,3-dione) (TheremoFisher) was diluted in DMSO to a concentration of 50 mM and used at a working concentration of 50 µM (1:1000 DMSO dilution). Both reagents were stored at – 80 °C. For analysis of percent change in response to drug application, 2 minutes of spontaneous activity after drug application was compared to the spontaneous activity after media or vehicle application.

### Immunocytochemistry

Cultures grown on coverslips were washed once with PBS, fixed with 4% paraformaldehyde for 10 minutes at room temperature, permeabilized with 0.1% Triton X-100 (Sigma) in a 3% bovine serum albumin (BSA) in PBS solution for 20 minutes at room temperature and washed three times with PBS before the addition of 3% BSA in PBS blocking solution for 60 minutes at room temperature. Primary (**Supplementary Table 2**) and secondary (**Supplementary Table 3**) antibodies were diluted in a 3% BSA in PBS and either 3% normal goat serum or 3% normal donkey serum (Millipore-Sigma) depending on the secondary antibody host species. The coverslips were incubated in primary antibodies for 16 hours at 4°C and washed three times with PBS before incubating in the secondary antibody solution for 1 hour at room temperature. Nuclear Stain Hoechst 33342 (Millipore Sigma) was added to the secondary solution at a dilution of 1:1000. Stained coverslips were mounted onto Superfrost Plus glass slides (ThermoFisher) using ProLong Antifade mounting media (ThermoFisher).

### Image acquisition, processing, and quantification

Low power images to quantify cell numbers across networks were performed on a Carl Zeiss Axio Imager.Z2 upright microscope equipped with a Carl Zeiss ApoTome.2 apotome using a 20X air objective. In cases where cultures on coverslips were analyzed quantitatively, the average of five frames was determined for each coverslip. Cell numbers were quantified using ImageJ software^36^.

3D confocal images were obtained on a Carl Zeiss LSM 880 Airyscan.2 microscope (Carl Zeiss, Oslo, Norway) with either a 63X or 25X oil immersion objective and analyzed using *Imaris* software (Oxford Instruments, Oxon, UK). Equal settings for laser intensity, step distance and pinhole aperture were used within individual experiments as much as possible.

Modifications to the brightness or contrast of images were performed equally to all images within an experiment. For Sholl analysis to ensure that excitatory neurons were being sampled, bipolar neurons were excluded and half of the neurons analyzed were counterstained with vesicular glutamate transporter 1 (VGLUT) and the statistics were not different between this group and the rest.

For synaptic quantification, *Imaris* software was used to generate 3D reconstructions of intracellular GFP and create a mask of proximal neurites. The pre-synaptic puncta were defined by vesicular glutamate transporter 1 (VGLUT1) or glutamic acid decarboxylase 65 (GAD65) immunoreactive puncta of 0.5 µm in diameter located outside of the neurite mask and within 1 µm of the external surface of the neurite. Post synaptic puncta were defined by post synaptic density 95 (PSD95) or GEPHRYN immunoreactive puncta of 0.3 µm within the neurite. Synapses were defined as pre- and post-synaptic puncta with 1 µm of each other. These were quantified and normalized to the area of the neurite determined by the GFP mask.

### Quantification and Statistical Analyses

Data are presented as the mean ± standard error of the mean (SEM). For single comparisons, two-way unpaired *t* tests were performed to determine significance at the level of *P* < 0.05. For multiple comparisons and grouped analysis, one way or two-way ANOVA with Tukey’s multiple comparison’s correction was performed. Sample sizes were not predetermined using statistical methods. Statistical analysis was performed using Prism 7 (GraphPad Software, Boston, MA, USA).

## Results

### 15q11.2 deletion alters network structure and cellular composition

HiPSC-derived cortical neurons were co-cultured with rat astrocytes beginning at day in culture (DIC) 37 and examined by florescence microscopy for markers of differentiation and maturation. By DIC 80, 6 weeks after co-cultures were established, the neurons derived from both control and 15q11.2 deletion bearing individuals exhibited features consistent with upper layer cortical neuron fate, as shown by the predominance of the post-mitotic, upper cortical layer transcription factor SATB2 (special AT-rich sequence-binding protein 2)^37^ (87.4 ± 4.3 % of neuron specific beta-tubulin III (TUJ1)^+^ neurons, n = 3 control donor lines vs 89.44 ± 0.7 %, n = 3 deletion donor lines; P = 0.67). There were also a small subset of neurons immunoreactive for SATB2 and the deep layer transcription factor CTIP2 (chicken ovalbumin upstream promoter transcription factor (COUP-TF)-interacting protein 2) ^37, 38^ with no significant differences between the two groups (10.5 ± 1.042 % TUJ1^+^ neurons, n = 3 control donor lines vs 14.26 ± 11.5 % n = 3 15q11.2 deletion donor lines; P = 0.76). At DIC 80, cultures consisted of dense TUJ1^+^ neurites connecting clusters of neuronal cell bodies (**Figure 1A**). However, there appeared to be fewer and more disorganized interconnecting neurites in the 15q11.2 deletion cultures (**Figure 1A**). This difference was more pronounced further in development at DIC 146 (**Figure 1B**), despite similar numbers of TUJ1^+^ neurons (26.9 ± 4.95 cells per high powered field, n = 4 control donor lines with 3 independent cultures per line vs 29.5 ± 4.64 cells per high powered field, n = 3 deletion donor lines with 2-3 independent cultures per line; P = 0.71). When neurons were stained for calcium calmodulin dependent kinase II (CAMKII) as an indicator of glutamatergic neuron fate^39–41^, most neurons were immunoreactive and there were similar proportions of CAMKII^+^ cells between the control and 15q11.2 deletion cultures at both DIC 80 (91.2 ± 1.2%, n = 4 control donor lines vs 93.1 ± 1.7%, n = 3 donor lines; P = 0.48) and DIC 146 (95.8 ± 0.9 vs 95.1 ± 1.4; P = 0.92) (**Figure 1A - D**). Immunofluorescent staining for glutamic acid decarboxylase 67 (GAD67), as an indicator of the amino acid gamma-aminobutyric acid (GABA) production^42, 43^, demonstrated a minority of cells staining positive in both control and 15q11.2 deletion lines. However, there was an increase in proportion of GAD67^+^ cells at DIC 80 (5.7 ± 0.9 %, N = 14 control cultures from n = 4 donor lines vs. 11.6 ± 1.4, N = 9 15q11.2 deletion cultures from n = 3 donor lines; P < 0.05) and a significant doubling of GAD67^+^ cells in the 15q11.2 deletion lines compared to controls at DIC 146 (7.1 ± 0.94 % N = 12 cultures from n = 4 control donor lines vs 18.0 ± 3.41 % N = 9 cultures from n = 3 15q11.2 deletion lines; P < 0.0001) (**Figure 1A - B, Figure 1E - F**). These data suggest alterations in cell fate and structural interconnectedness in 15q11.2 deletion neurons.

**Figure 1.**
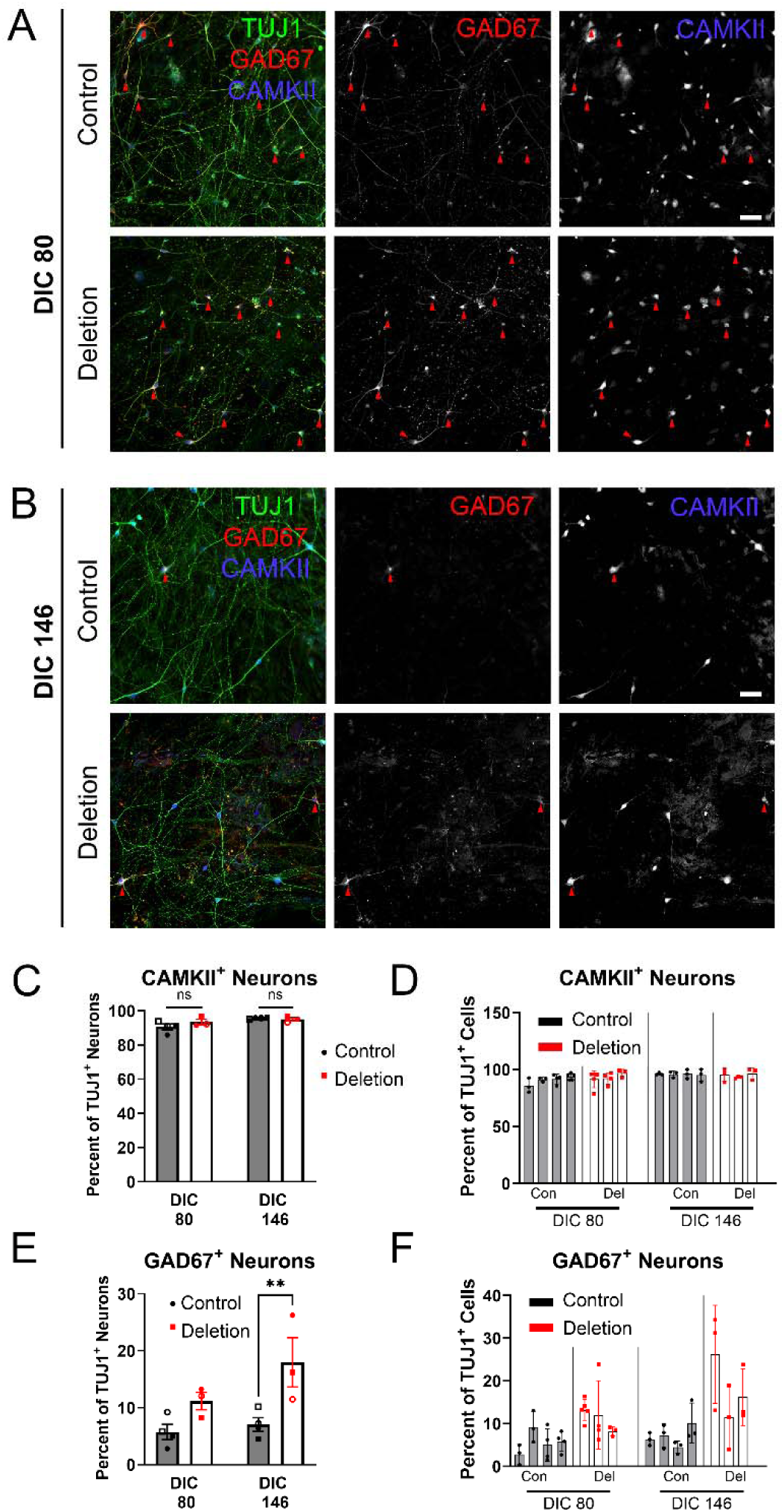
15Q11.2 deletion alters network structure. (A-B) Representative images of TUJ1^+^ (green, left merged panel) neurons stained for CAMKII (blue) and GAD67 (red) in control and deletion neurons to identify glutamatergic and GABAergic neurons, respectively demonstrate deficits in network organization. Images taken on day in culture (DIC) 80 (A) and DIC 146 (B). Red arrows indicate GAD67^+^ neurons. Scale bar 50 µm. (C) Quantification of the percent of TUJ1^+^ neurons expressing CAMKII at DIC 80 and 146. Each symbol represents the mean of a minimum of three cultures from an individual donor line. Bars are the mean ± standard error of the mean (SEM) of four control and three 15q11.2 deletion lines. (D) Data from C presented by donor line with each column representing a donor and each symbol representing an independent culture. (E) Quantification of the percent of TUJ1^+^ neurons expressing GAD67 at DIC 80 and 146. Each symbol represents the mean of a minimum of three cultures from an individual donor line. Bars are the mean ± standard error of the mean (SEM) of four control and three 15q11.2 deletion lines. Two-way ANOVA with Tukey’s multiple comparison. ** = *P* < 0.01. (F) Data from D presented by donor line as in D.

### Decreased neurite outgrowth and complexity in 15q11.2 deleted neurons

The 15q11.2 region contains at least one gene, CYFIP1, that encodes a protein involved in both cytoskeletal and synaptic development. The observation that cultures are less interconnected in the 15q11.2 deletion cultures (**Figure 1A**) suggests that there may be primary deficits in neurite growth and expansion. To examine cytoskeletal changes, hiPSC-derived cortical forebrain neurons in co-culture with rat astrocytes were sparsely transfected with a plasmid expressing the green fluorescent protein (GFP) under the promotor for SYNAPSIN (pAAV-hSYN-EGFP) (SYNAPSIN-GFP) and matured until DIC 80 and 146 for single cell neurite analysis.

The morphology of the neurons from 15q11.2 deletion lines was less complex with an fewer neurites and branches when compared to control lines at both DIC 80 and 146 (**Figure 2A-B**). Sholl analysis of DIC 80 neurons demonstrated a significant reduction in the number of intersections from 20 to 160 μm from the soma between deletion and control lines (**Figure 2A and 2C**). At DIC 146 there was also a significant difference in the number of intersections between control and deletion lines at 30 µm and from 60 µm to 180 µm (**Figure 2B and 2D**). As expected for expanding neuronal arbors, the distance to the peak number of intersections increased in both populations over time. This distance to peak and the value of the peak number of intersections was greater in control compared to 15q11.2 deletion neurons at both timepoints (**Figure 2C and D**). At 80 µm, the distance at which the peak number of intersections occurred in control neurons at day 146, there were significantly fewer intersections in the 15q11.2 deletion cells at DIC 80 compared to controls (control = 5.17 ± 0.119 intersections, N = 29 neurons form n = 3 donor lines vs 15q11.2 deletion = 2.73 ± 0.253 intersections, N = 25 neurons from n = 3 donor lines; P = 0.039). At DIC 146 there continued to be fewer intersections in 15q11.2 deletion neurons (5.9 ± 0.8, N = 45 neurons from n = 3 donor lines) compared to the control neurons (8.4 ± 0.51, N= 49 neurons from n = 4 donor lines; P = 0.0239) (**Figure 2E-F**). While the number of intersections increased from DIC 80 to 146, the number of intersections in the deletion bearing cells at day 146 was only equal to the number in controls at DIC 80 (P = 0.758), suggesting a delay in structural maturation.

**Figure 2.**
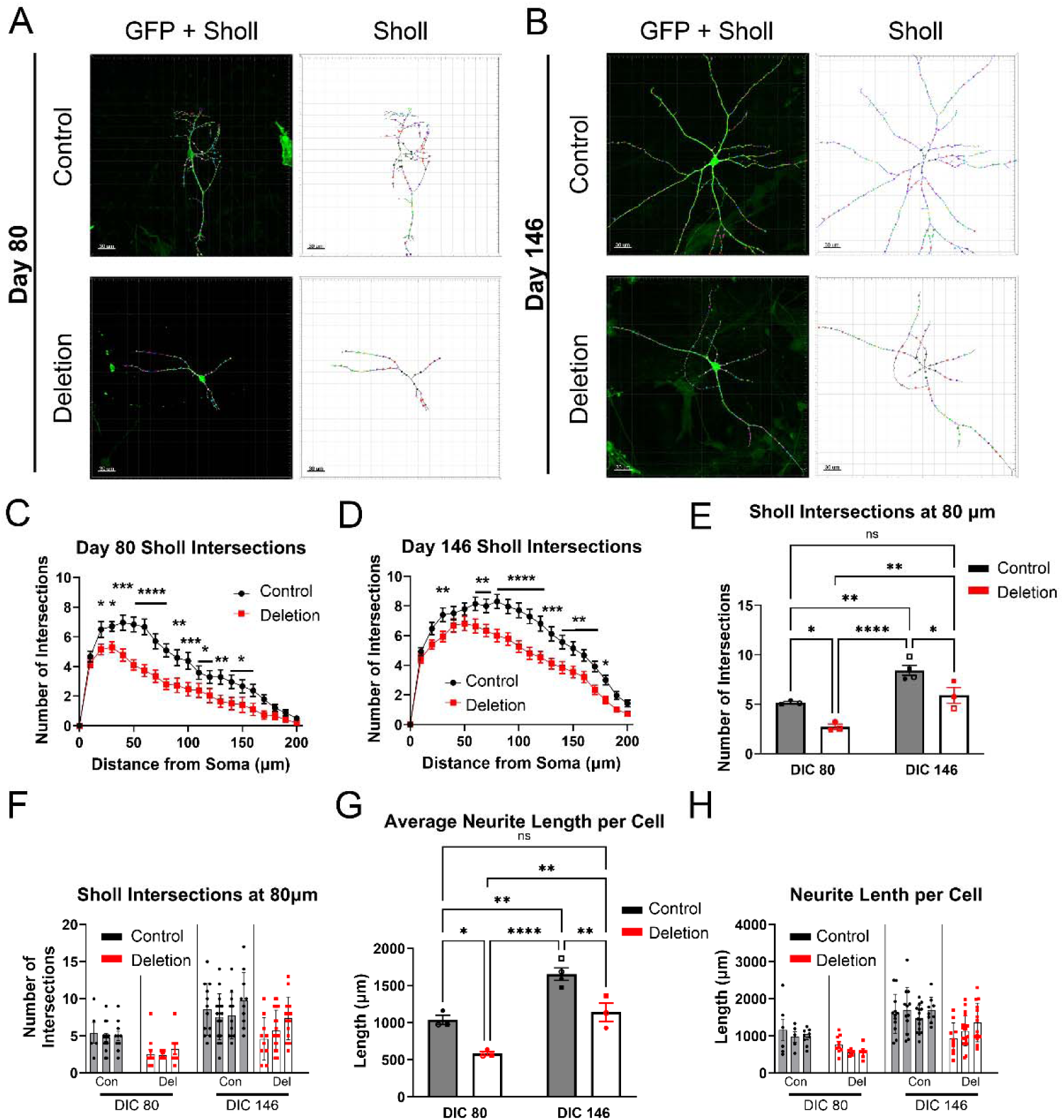
15q11.2 deletion decreases dendritic complexity and length. (A-B) 3D reconstructions of 25X oil immersion confocal images of *pAAV-hSyn-EGFP* (SYNAPSYN-GFP) expressing representative neurons with corresponding Sholl intersections in control and 15q11.2deletion neurons at DIC 80 (A) and DIC 146 (B). Panels on the left demonstrate reconstruction of GFP (green) with Sholl intersection overlay (multicolored spheres). Panels on the right demonstrate inverted images with Sholl intersection and neurite skeleton only. Scale bar 50 µm. (C) Quantification of Sholl analysis of N = 29 control cells from 4 lines and N= 26 deletion cells from 3 lines at DIC 80. Two-way ANOVA *P* < 0.0001. (D) Sholl analysis of N = 49 control cells from 4 lines and N = 45 deletion cells from 3 lines at DIC 146. Two-way ANOVA *P* < 0.0001. (E) Average Scholl intersections by line at 80 µm from data in B and C. Individual data points are means of 6 to 12 cells per donor line. Columns represent means ± SEM of n = three to four donor lines. Individual symbols are the mean of a single donor line. Two-way ANOVA *P* < 0.001. (F) Data from E presented by donor line. Columns represent mean per line and symbols are individual neurons. (G) Average total neurite length per cell is decreased in deletion compared to control donor lines. Symbols are average of cells from each donor line. Columns are means of N = 3 or 4 donor lines per condition and timepoint. (H) Data from G presented per donor line. Column is the mean per line and each symbol is a neuron. Two-way ANOVA *P* < 0.001. Tukey’s multiple comparisons * = *P* < 0.05, ** = *P* < 0.01, *** = *P* < 0.001, **** = P < 0.0001.

In addition to fewer intersections, the total dendritic length per cell was decreased in the 15q11.2 deletion neurons at both DIC 80 (Mean difference 453 µm; 95% CI 68.97 to 837.1; P = 0.0216) and DIC 146 (513.5 µm; 68.97 to 837.1; P = 0.007), with the deletion neurons achieving the same length at DIC 146 as the controls at DIC 80 (-102.3 µm; -486.4 to 281.8; P = 0.8384) (**Figure 2G-H**). Together with the differences in complexity of the arbors, the difference in total dendritic length indicates a delay in structural maturation in the 15q11.2 deletion lines.

### Proximal synaptic density is not affected by 15q11.2 deletion

The simplicity of the neurite tree revealed by the experiments in **Figure 2** could be due to a primary cytoskeletal, adhesion or synaptic defect. Early changes in neurite outgrowth argue for a primary cytoskeletal or adhesion defect, while the increased magnitude of differences later in development argue that synaptic defects contribute to these changes. We examined the density of both glutamatergic and GABAergic synapses onto the proximal neurites of SYNAPSIN-GFP transfected cells at DIC 146 (**Figure 3**). Excitatory synapses were assessed by 3D confocal microscopy of co-cultures immunolabeled for presynaptic VGLUT and post synaptic PSD95(**Figure 3A**). The resultant synapse density was similar between control and 15q11.2 deletion cultures (Two-tailed unpaired t-test, P = 0.19) (**Figure 3B**).

**Figure 3.**
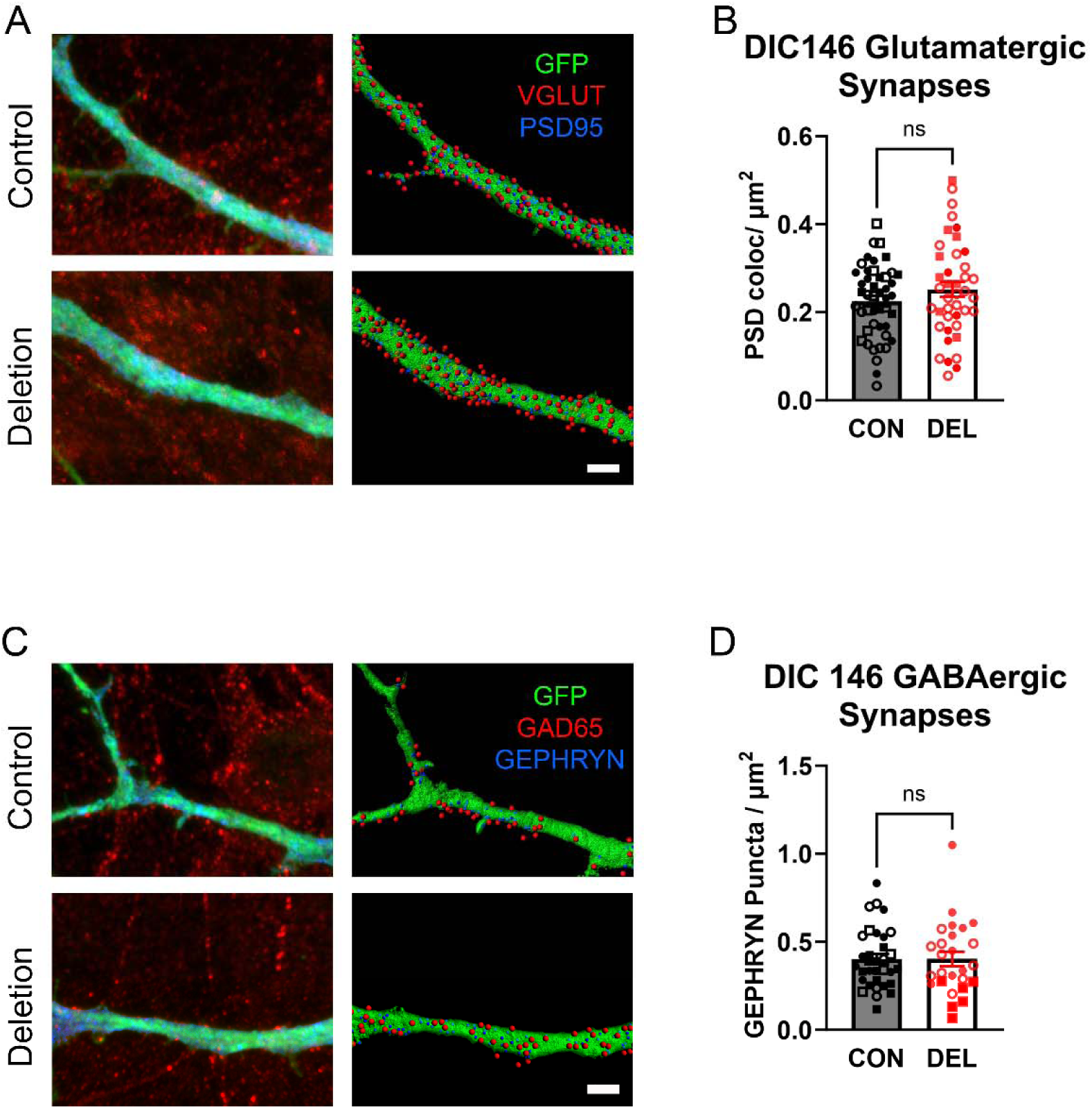
Proximal synaptic density in 15q11.2 deleted neurons. (A) Representative 3d reconstructions of 63X confocal images of control (top) and 15q11.2 deletion (bottom) proximal neurites of SYNAPSIN-GFP expressing neurons at DIC146 stained for GFP (green), vesicular glutamate transporter 1 (VGLUT1) (red) and post synaptic density protein 95 (PSD95) (blue) are shown in left panels. Panels on the right show the mask generated by GFP staining (green). The presynaptic puncta from glutamatergic synapses on to the masked neurons are generated from VGLUT1 staining outside of the mask and within 1 µm of the neurite are in red. Post synaptic puncta re in blue and are generated from PSD95 staining within the masked region. (B) Quantification of glutamatergic synapses, defined as pre and post synaptic puncta colocalizing within 1 µm of each other, normalized to the surface area of the masked region to determine density. Each symbol type represents a different donor line with each symbol representing a neurite from a unique cell. (C) Left panels show representative 3d reconstructions of 63X confocal images of control (top) and 15q11.2 deletion (bottom) proximal neurites of SYNAPSIN-GFP expressing neurons at DIC146 stained for GFP (green), GAD65 (red) and post synaptic scaffolding protein GEPHRYN (blue). Panels on the right show the mask generated by GFP staining (green). The presynaptic puncta from GABAergic synapses on to the masked neurons are generated from GAD67 staining outside of the mask and within 1 µm of the neurite. Post synaptic puncta are generated from GEPHRYN staining within the masked region. (D) Quantification of glutamatergic synapses, defined as pre and post synaptic puncta colocalizing within 1 µm of each other, normalized to the surface area of the masked region to determine density. Each symbol type represents a different donor line with each symbol representing a neurite from a unique cell. Scale bar 3 µm. Not significant by Two-tailed t test.

Inhibitory synapses were labeled with antibodies targeting presynaptic GAD65 and the inhibitory post synaptic scaffolding protein, GEPHRYN (**Figure 3C**). There was no significant difference between the control and the 15q11.2 deletion cultures (Two-tailed unpaired t test, P = 0.9917) (**Figure 3D**). These findings suggest that the 15q11.2 deletion does not determine the density of glutamatergic or GABAergic synapses, but do not speak to functional or structural changes within individual synapses. Additionally, with the same density of synapses, the absolute number of synapses would be decreased with a smaller neurite arbor in the 15q11.2 deletion neurons and does not preclude a functional change at the network level.

### 15q11.2 deletion neurons display delayed functional maturation

The apparent deficits in neurite development and altered network morphological interconnectivity suggests that functional neuronal development may be affected by the 15q11.2 deletion. To assess the effects of 15q11.2 deletion on functional maturation of neurons, co-cultures of hiPSC derived cortical neurons and rat astrocytes were plated onto 24-well MEA plates at DIC 37. Voltage recordings were conducted beginning at DIC 39 and subsequently every one to two weeks until DIC 202. Discrete spikes recorded from each electrode over time (**Figure 4A**,) were used to determine weighted mean firing rate (WMFR), burst firing and eventually synchronized network bursts. Spikes appeared in some cultures as early as DIC 39, and consistent spiking across multiple electrodes were recorded after day 50 (Figure **4A-B**), with consistent busting after DIC 60 (**Figure 4A and 4C**) and synchronized network bursting after DIC 70 (**Figure 4A and 4D**).

**Figure 4.**
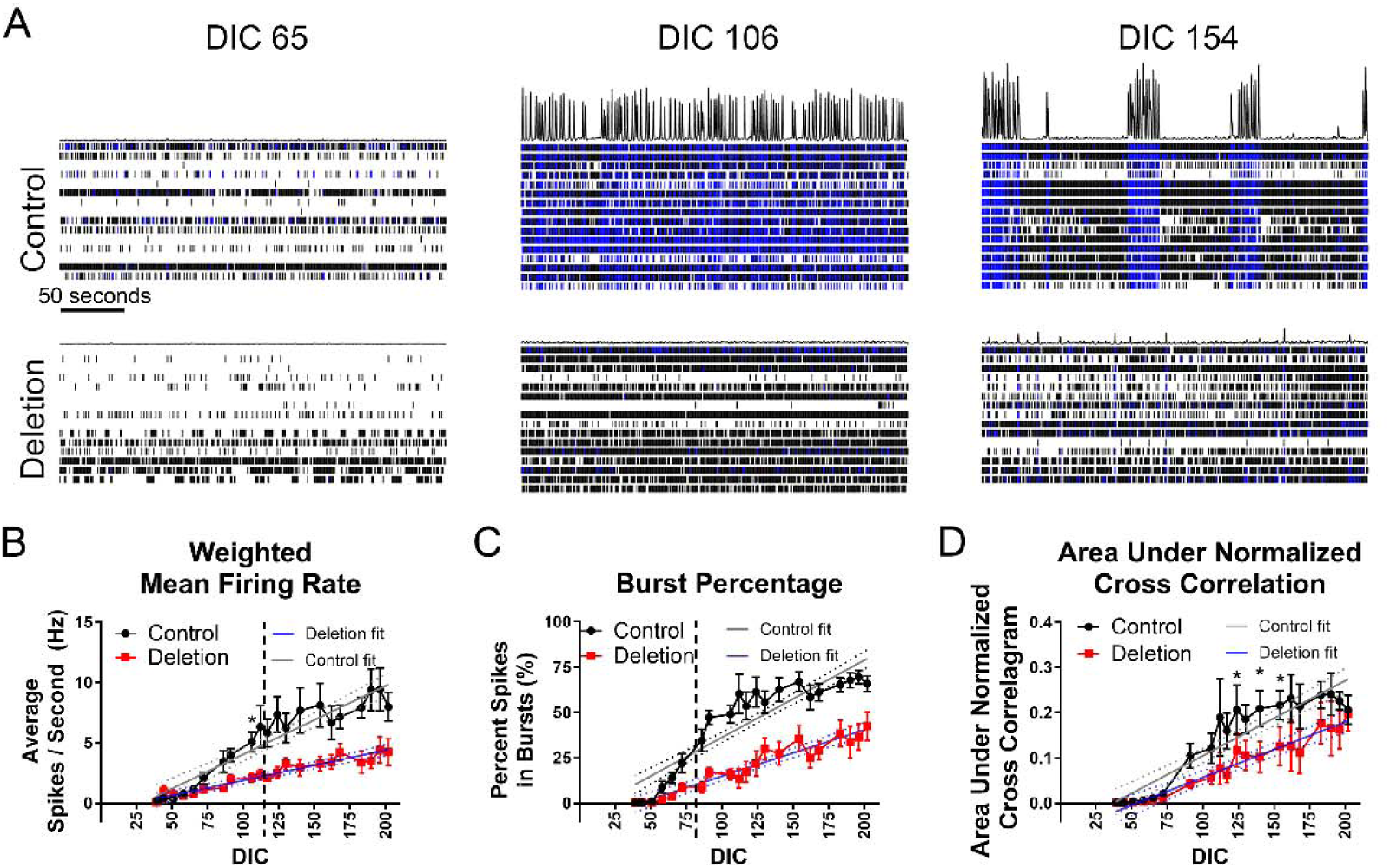
15q11.2 deletion decreases firing rate and bursting and delays synchrony in culture networks. (A) Raster plots of spikes determined from voltage recordings from 16 electrodes from representative single wells of control (top) and 15q11.2 deletion (bottom) cultures at DIC 65, 106 and 154. Each row is data from a single electrode. Single spikes for a given electrode are represented by black vertical marks. Blue vertical lines are spikes on a single electrode meeting criterion for an electrode burst (5 spikes occurring within 100 ms). Black tracing at the top of each panel represents relative spikes per time per second across the entire well. Time scale bar is 50 seconds. Analysis of four control and three 15q11.2 deletion lines over at least 3 independent differentiations for (B) weighted mean firing rate (Two-way ANOVA P < 0.0001), (C) percent of spikes that occur in bursts (burst percentage) (Two-way ANOVA *P* <0.0001), Tukey’s multiple comparison test and (D) area under normalized cross correlogram (AUNCC) as a quantification of synchronization (Two Way ANOVA *P* < 0.0001). Mean values ± SEM are indicated by symbols. Linear regressions (solid lines) with 95% confidence intervals (dashed lines) are in grey and blue. Two-way ANOVA *P* < 0.0001. Tukey’s multiple comparisons * = *P* < 0.05, ** = *P* < 0.01, *** = *P* < 0.001, **** = P < 0.0001. For C and D, Tukey’s multiple comparison’s testing indicates significance from ** to **** for all data points to right of the vertical dashed line.

In 15q11.2 deletion cultures, while spikes emerged at similar time points as controls, the weighted mean firing rates (WMFR) showed delayed maturity with significantly different slopes (Control slope = 0.057, 95% CI 0.047 to 0.067 vs Deleted slope = 0.025, 95% CI 0.020 to 0.030; P < 0.0001) and significant differences between the two populations at DIC 106 (P < 0.05) and at all time points From DIC 117 to DIC 202 (P < 0.05 to P < 0.001) (**Figure 4B**). Bursting emerged consistently in control neurons at DIC 58 and there was a progressive increase in the percentage of spikes that occurred in bursts until DIC 130 when this plateaued between 50 and 75%. In contrast, the 15q11.2 deletion cultures exhibited less bursting at DIC 58, and the increase in burst percentage was lower and continued to increase at the point the control cultures had leveled off, with a significant difference between control and deletion at DIC 86 and at every timepoint afterwards (P < 0.05 to P < 0.0001) (**Figure 4C**).

As cultures mature, neurons begin to exhibit synchronized periods of high activity alternating with low activity across most electrodes (**Figure 4A**). These spontaneous slow network oscillations started by DIC 70 in control cultures, consistent with the development of functional synaptic connectivity between neurons and synchronization of the network ^44^. To assess for the development of synaptic synchronization of the network across time, the area under normalized cross correlation (AUNCC) from each culture and at each time point was determined. The AUNCC is a measure of the area under interelectrode cross-correlation normalized to the autocorrelation, and is a measure or synchronicity of the network with higher values indicating greater synchronicity ^35^. In controls, synchronized activity occurred later than bursting, with a significant rise between DIC 65 and 106 (**Figure 4A, and Figure 4D**). However, the development of synchronized activity occurred over a more protracted time course in the 15q11.2 deletion neurons (Control slope = 0.057, 95% CI 0.0014 to 0.0020 vs Deleted slope = 0.0012, 95% CI 0.00096 to 0.0015; P <0.05) (**Figure 4D**). This suggests altered functional development of synaptic connectivity as well as basic functional neuronal properties.

### 15q11.2 hiPSC-neurons demonstrate a delay in AMPA / kainite receptor-mediated signaling

To determine whether differences in synaptic activity contribute to differences in the functional maturation of these nascent neuronal networks, we examined glutamatergic network connectivity by blocking the α-amino-3-hydroxy-5-methyl-4-isoxazolepropionic acid (AMPA) and kainate glutamate receptors with CNQX. We examined three timepoints at which robust spiking and bursting had already been established in control cultures (**Figure 5**). As might be expected for early network development when there is less or variable synaptic activity, the effects of 50 µM CNQX application at the earliest timepoint, DIC 100, were equivalent in both the control and deletion cultures with similar numbers of cultures demonstrating an increase or a decrease in spiking, bursting or synchronized activity (**Figure 5A-D**). There was no significant difference in the change in WMFR between control and 15q11.2 deletion cultures at any timepoint even though CNQX had a progressively larger effect on firing over time (**Figure 5B**). However, at DIC 134, most control cultures responded to CNQX with a marked reduction in burst percentage and there was a significant difference between control and deletion cultures with less of a decrease in deletion cultures (control = -46.64 ± 7.7 %, n = 25 cultures; vs. 1511.2 deletion = -7.7 ± 18.7 %, n = 20 cultures; P = 0.012). By DIC 210, the deletion cultures had a similar reduction in burst percentage; suggesting a developmental delay that is overcome (Control = -61.261 ± 4.049 %, n = 15; vs. 1511.2 deletion = -60.7 ± 9.2 %, n = 12; P = 0.98) (**Figure 5C**).

**Figure 5.**
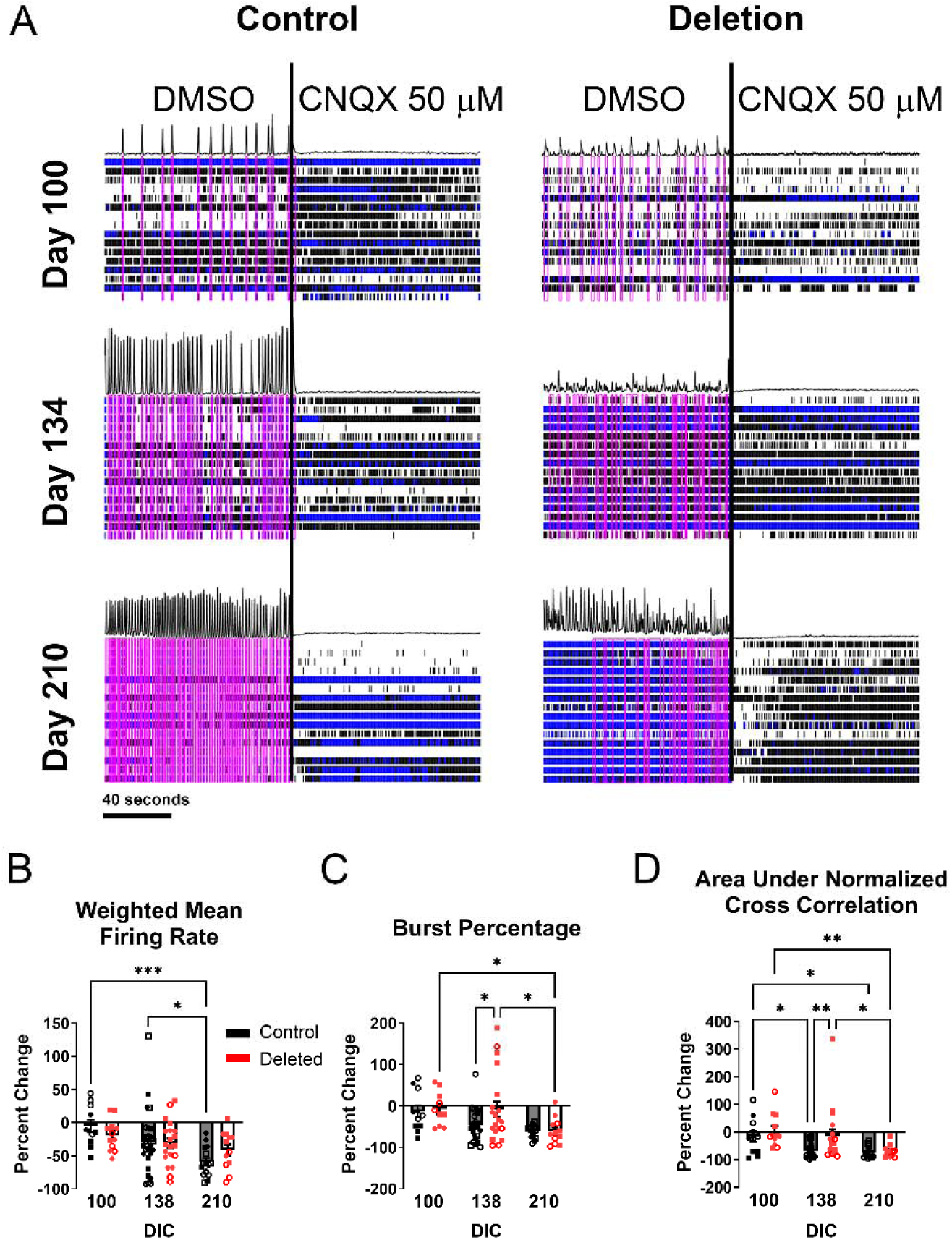
15q11.2 deletion results in delayed glutamatergic influence on the network. (A) Raster plots of spikes determined from voltage recordings from 16 electrodes from representative single wells of control (left) and 15q11.2 deletion (right) cultures. Each row is data from a single electrode. Single spikes for a given electrode are represented by black vertical marks. Blue vertical lines are spikes on a single electrode, meeting criteria for an electrode burst - 5 spikes occurring within 100 ms. Black tracing at the top of each panel represents relative spikes per time per second across the entire well. Pink boxes are network bursts. Time scale bar is 40 seconds. Black vertical line demarcates time of acute CNQX application. Quantification of CNQX mediated effects on(B) weighted mean firing rate (WMFR), (C) burst percentage and (D) synchronization in control and deletion cultures. Each data point is the mean activity across 16 electrodes for a single well. Individual symbols are independent cultures with the type of symbol indicating a unique donor. Columns are the mean ± SEM of all cultures with 4 control and 3 deletion culture represented. Two-way ANOVA *P* <0.0001. Tukey’s multiple comparisons * = *P* < 0.05, ** = *P* < 0.01, *** = *P* < 0.001.

Like developmental changes in burst percentage, CNQX-dependent changes in synchronization were slower to emerge in the 15q11.2 deletion cultures, with a significantly smaller decrease in synchronicity in response to CNQX at DIC 134 (Control = -69.3 ± 5.3 %, n = 25; vs. 1511.2 deletion = -34.316 ± 12.107 %, n = 17; P = 0.01). By DIC 210, the deletion cultures have similar reductions in synchronicity in response to CNQX as the control cultures (Control = -73.494 ± 6.084 %, n = 14; vs. 1511.2 deletion = -67.0 ± 6.6 %, n = 11, P = 0.71) (**Figure 5D**). Taken together, these data suggest that 15q11.2 iPSC-neurons have a delay in the development of AMPA / Kainate receptor-mediated synaptic signaling.

### Development of an inhibitory GABAergic response is altered in 15q11.2 deletion neurons

We evaluated the effects of GABAergic influence on network development using the GABA-A receptor antagonist bicuculline. At DIC 79 and DIC 124, application of bicuculline decreased the WMFR in both control and 15q11.2 deletion cultures (**Figure 6A and B**). A decrease in firing in response to blocking GABA-A receptors suggests that the overall effect of baseline endogenous GABAergic transmission on the network is increased neuronal firing, consistent with the presence of excitatory GABAergic signaling in immature neurons ^45^. There was a non-significant reduction in the amount of bicuculline-dependent inhibition between DIC 79 and 124 in both deletion and control cultures. However, by DIC 148 in the control cultures, there was a switch in the response to bicuculline to one that reflects an increase in firing rate (127% ± 26.2%; n = 11) while the response in 15q11.2 deletion cultures is still a decrease in firing (-15.9% ± 12.02%; n = 12) (P < 0.0001) (**Figure 6B**). Analysis of burst percentage in response to bicuculline also revealed no significant difference in the mean percentage change between control and 15q11.2 deletion cultures and either a reduction in firing or an equivocal response at DIC 79 and 124. However, at DIC 148 all control cultures demonstrated an increase in burst percentage in response to bicuculline (56.5 ± 25.6 %, n = 10) while15q11.2 deletion cultures continued to demonstrate a reduction in burst percentage in response to bicuculline (- 51.9 ± 10.3%, n = 11) (P = 0.012) (**Figure 6A and C**). Similarly, the bicuculline dependent change in the AUNCC was either equivocal or negative in both the control and deletion cultures at DIC 79 and 124, but the two genotypes diverged at DIC148 with an increase in synchronicity based on a positive percent change in control cultures (93.4 ± 20.4 % n = 11) and an equivocal response in the 15q11.2 deletion cultures (-2.3 ± 19.0 %, n = 11) (P = 0.006) (**Figure 6D**). These findings suggest either a delay in the maturation of a chloride gradient or a delay in synaptic GABAergic activity in the 15q11.2 deletion cultures and are consistent with overall structural and functional delays.

**Figure 6.**
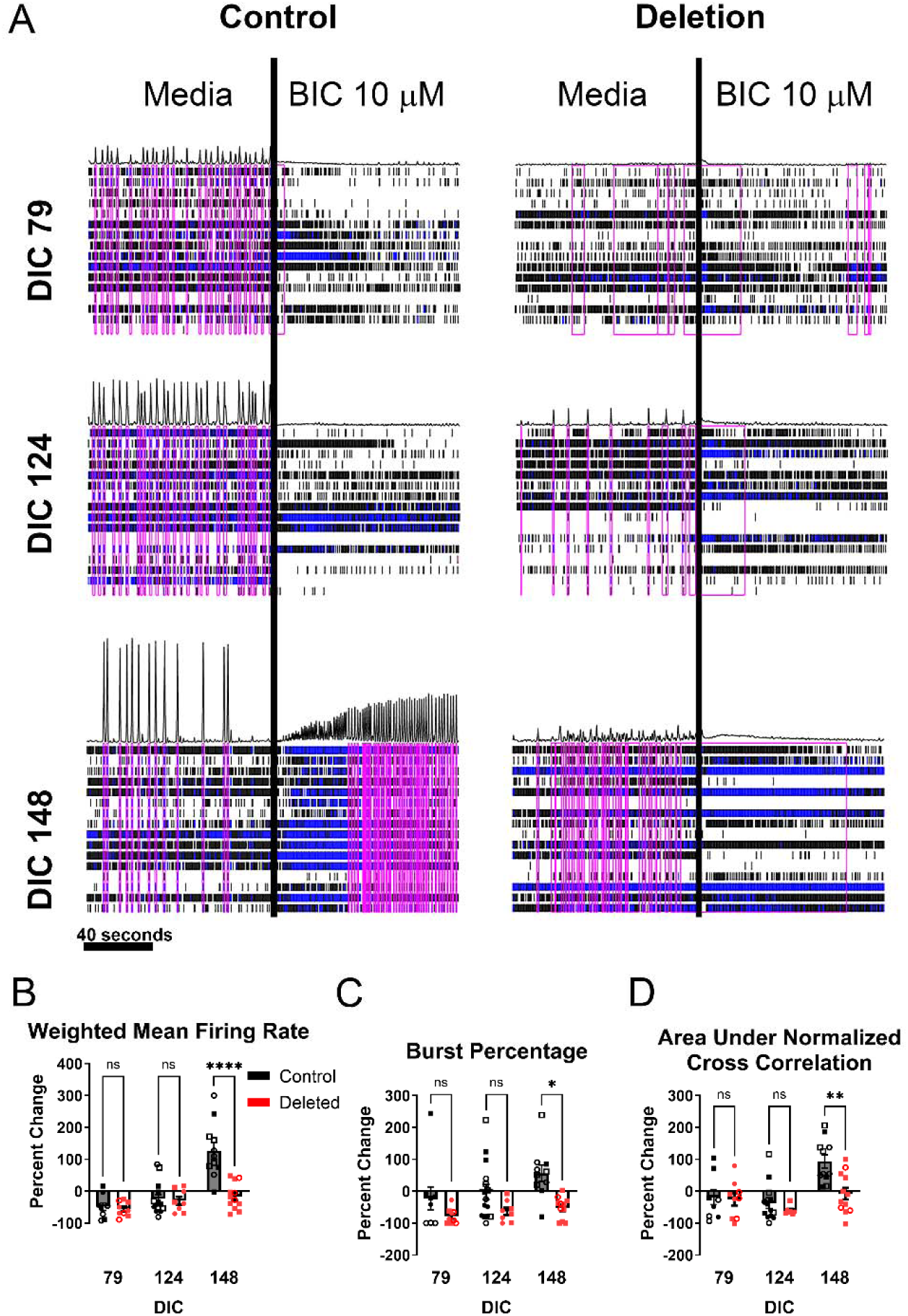
15q11.2 deletion delays the maturation of synaptic GABA responses. (A) Example raster plots demonstrating acute GABA receptor blockade with application of bicuculline. Raster plots of spikes determined from voltage recordings from 16 electrodes from representative single wells of control (left) and 15q11.2 deletion (right) cultures. Each row is data from a single electrode. Single spikes for a given electrode are represented by black vertical marks. Blue vertical lines are spikes on a single electrode meeting criterion for an electrode burst - 5 spikes occurring within 100 ms. Black tracing at the top of each panel represents relative spikes per time per second across the entire well. Pink boxes are network bursts. Time scale bar is 40 seconds. Black vertical line demarcates time of acute Bicuculline application. Quantification of bicuculline mediated effects on(B) weighted mean firing rate (WMFR), (C) burst percentage and (D) synchronization in control and deletion cultures. Each data point is the mean activity across 16 electrodes for a single well. Individual symbols are independent cultures with the type of symbol indicating a unique donor. Columns are the mean ± SEM of all cultures with 4 control and 2 to 3 deletion culture represented. Two-way ANOVA. Tukey’s multiple comparisons * = p < 0.05, **= p < 0.01, *** = p < 0.001, **** = p < 0.0001.

## Discussion

While 15q11.2 deletion has been shown to alter NPC biology^17^, this is the first detailed assessment of later neuronal development and function in a human cell model. This study demonstrates that deletion of the 15q11.2 region is sufficient to cause structural changes at the cellular level including decreased neurite length and complexity. This is associated with early altered structural connectivity at the network level and functional delays in spontaneous activity and later deficits in synaptic mediated firing, bursting and synchronization. This developmental dysregulation *in vitro* suggests a possible cellular mechanism for the phenotypes of individuals affected by the 15q11.2 deletion, namely that deficits in neurite development alter maturation of both glutamatergic and GABAergic synaptic connectivity.

The overall decrease in spiking, spike bursting and synchronized activity in 15q11.2 deletion neurons that is shown in this study to be due in part to a functional delay in AMPA / Kainate receptor mediated connectivity is consistent with postmortem studies of patients with schizophrenia having decreased dendritic spines and other iPSC models of schizophrenia indicating impaired glutamatergic transmission ^46^. However, at least in the proximal portions of the neurite arbor this was not due to a significant change in the density of excitatory synapses. There may be structural and functional changes within the synapses that cannot be identified at the resolution used here or changes in density more distally. Regardless of any additional functional or structural synaptic changes, the overall decreased neurite length and arborization results in overall decreased excitatory input despite similar synaptic densities.

Given the predominance of epilepsy phenotypes in 15q11.2 deletion bearing individuals, the decreased activity and glutamatergic influence of neuronal networks, *in vitro,* is somewhat counterintuitive to the prevailing hypothesis that the underlying cause of epilepsy is hyperexcitability, in part due to increased excitatory activity ^47^. However, loss of appropriate inhibition can also lead to pathologic hyperexcitable states. In this study, there is a delayed maturation of the inhibitory response to GABA, demonstrated by application of bicuculline. Immature neurons in brain development have higher intracellular chloride than mature cells and the activation of GABA receptors results in chloride efflux and neuronal depolarization. GABA is excitatory early in development, is required for primitive network activity, regulates proliferation and migration and strongly influences synaptic plasticity ^48^. A prolonged window of excitatory GABAergic activity or failure to establish the appropriate chloride gradient can pathologically alter the excitation inhibition balance and synaptic plasticity through numerous mechanisms. Cultures of 15q11.2 deletion neurons not only demonstrate a continued excitatory response to GABA at time points where the control neurons are inhibited, but there is an increase in the percentage of neurons in the culture that express GAD67, indicative of GABAergic neuron maturation. The presence of increased numbers of neurons expressing both GAD67 in the 15q11.2 deletion cultures could indicate a primary difference in developmental fate choice or a reactive increase in response to dysfunctional GABAergic signaling, but in the setting of prolonged excitatory GABA, offers another possibility for altered balance.

Altered dendritic development and synaptic function predispose individuals to neurodevelopmental disorders such as schizophrenia, epilepsy and ID. In schizophrenia, changes in dendritic spine density and morphology and decreased dendritic arborization, length and number have been shown in the prefrontal cortex^49, 50^. Decreased dendritic arborization has not only been shown in postmortem studies but also in living patients with neurite density imaging modeling of MRI data ^51, 52^ and is consistent with the changes demonstrated in this study, *in vitro,* using 15q11.2 deletion neurons.

Variable changes in dendrite and dendritic spine density have also been described in multiple types of epilepsy. In type II focal cortical dysplasia, synaptic spine density is decreased on the dendrites of dysplastic neurons and dendritic branching is altered with increased and disorganized dendrites in dysplastic neurons and decreased dendritic branch points in the relatively normal adjacent neurons^53^. Dendritic branch points, dendritic length and dendritic spines are also decreased in surgical samples from humans with temporal lobe epilepsy ^54–57^. Similar findings of decreased branch points and dendritic lengths have been described in syndromic forms of intellectual disability with an increased risk of epilepsy in Down syndrome and Rhett syndrome ^58^. However, particularly when looking at post-surgical or postmortem cases of epilepsy and schizophrenia, the question of whether the changes are causative or reactive to years of abnormal excitation and medication effects lingers. The results of the current study suggest that in the case of 15q11.2 deletion, a primary defect in dendrite development that results in a functional change in the activity of neuronal networks is a common mechanism for the different phenotypes.

Disease causing copy number variants involving multiple genes are difficult to study because manipulation of one gene may not be sufficient to recapitulate a phenotype. hiPSC derived cells offer the ability to study human cells with the genetic change directly but what is known about single genes in these regions can also provide significant insight. There are four genes that located at the 15q11.2 BP1-BP2 locus. *NIPA1* is expressed in neuronal tissue and mediates Mg^++^ transport as well as transport of other divalent cations. NIPA1 Pathogenic variants, but not deletions result in autosomal dominant hereditary spastic paraplegia (HSP) ^59^. Interestingly, although initially described as a pure phenotype of spastic paraplegia, families with HSP due to certain pathogenic variants in NIPA1 have also been reported to have epilepsy and ALS possibly related to reduced protein expression ^60, 61^. NIPA2 is a selective magnesium transporter that mediates renal magnesium conservation ^62^. TUBGCP5 is a member of a family of 5 similar proteins that bind gamma tubulin and function in spindle assembly ^63, 64^. While *NIPA1* and *TUBGCP5* have the potential to be important in brain physiology, *CYFIP1* is proposed to be a critical regulator of neural development and synaptic plasticity ^22, 26^.

CYFIP1 is highly conserved with expression in both excitatory and inhibitory neurons and complete knockout in mice is embryonic lethal ^21, 26^. Down regulation of the CYFIP1 protein leads to disruption of the organization of NPC’s in embryonic mice *in vivo* ^17^ as well as altered proliferation of pre- and postnatal neural progenitor cells^18^, suggesting that it plays an important role in early neural development. Haploinsufficiency in mice leads to decreased dendritic complexity^65^ similar to what is demonstrated here in 15q11.2 deletion neurons. The most likely mechanism for CYFIP1 mediated changes in dendritic structure is dysregulation of the WAVE regulatory complex (WRC) which is required for actin nucleation in dendritic branching and elongation ^66^. Studies in mice with reduced CYFIP1as well as in human cells with 15q11.2 deletion cells have demonstrated that a reduction of CYFIP1 results in a reduction of multiple other WRC proteins ^67^. Haploinsufficiency of *Cyfip1* in rodents also results in immature synaptic spine structure and function due to its dual interactions with FMRP and the WAVE regulatory complex ^65, 68, 69^. It is present at both glutamatergic and GABAergic synapses and decreased expression has been shown to stabilize GABAergic synapses in mature neurons ^24^. While this study did not determine a difference in gross density of synapses as a result of 15q11.2 deletion, electrophysiology demonstrated delays in both synaptic systems. This could be a result of overall reduced connectivity due to decreased numbers and length of dendrites or CYFIP1 mediated molecular changes at the synapses themselves.

## Conclusions

15q11.2 deletion results in decreased neurite complexity at the single cell level. At the network level, there is altered structural interconnectedness, proportions of glutamatergic and GABAergic neurons, and delays in the maturation of spike rates, busting and synchronization at the network level. This is associated with a decrease in the influence of AMPA/ Kainate receptors on the network (glutamatergic synaptic excitation) as well as a prolonged excitatory response to GABA. These findings suggest several mechanisms by which neuronal network deficits can lead to pathological changes in human development and result in the complex phenotypes manifesting in 15q11.2 deletion bearing individuals, but interpretation is limited by the simplified networks present *in vitro*. Some, but not all the neuronal changes in 15q11.2 deletion neurons are like what has been described in animal models of *Cyfip1* deletion and further exploration of the intersection between *in vivo* animal models and *in vitro* human models is likely to shed light on pathologic development in this disorder as well as potential therapeutic strategies.

## Acknowledgements

CNCDPK12, K08NS102526, Doris Duke Foundation Clinical Scientist Award to CWH; NIH 5R01NS117604 and Department of Defense ALSRP W81XWH1810175 to N.J.M.; NIH R35NS116843 to H.S. and NIH R35NS097370 to G-l.M. Department of Department of Neuroscience Imaging center, Johns Hopkins University. Haiwen Chen, MD PhD and Sean Oddoye, BS for logistical support.

---

**Supplementary Table 1:**

| Cell Line | Type | Sex | Age | Source |
| --- | --- | --- | --- | --- |
| C1.2 | Control | Male | fetal | Coriell<br>Biorepositories |
| C3.1 | Control | Female | 40 | Coriell<br>Biorepositories |
| GM01582 | Control | Female | 11 | Coriell<br>Biorepositories |
| CS8PAA | Control | Female | 58 | Cedar Sinai |
| CS0002 | Control | Male | 51 | Cedar Sinai |
| CS9XH7 | Control | Male | 53 | Cedar Sinai |
| Y1.3 | Deletion | Male | 52 | Coriell<br>Biorepositories |
| Y3.2 | Deletion | Female | 14.8 | Coriell<br>Biorepositories |
| Y4.3 | Deletion | Female | 46.4 | Coriell<br>Biorepositories |

**Supplementary Table 2:**
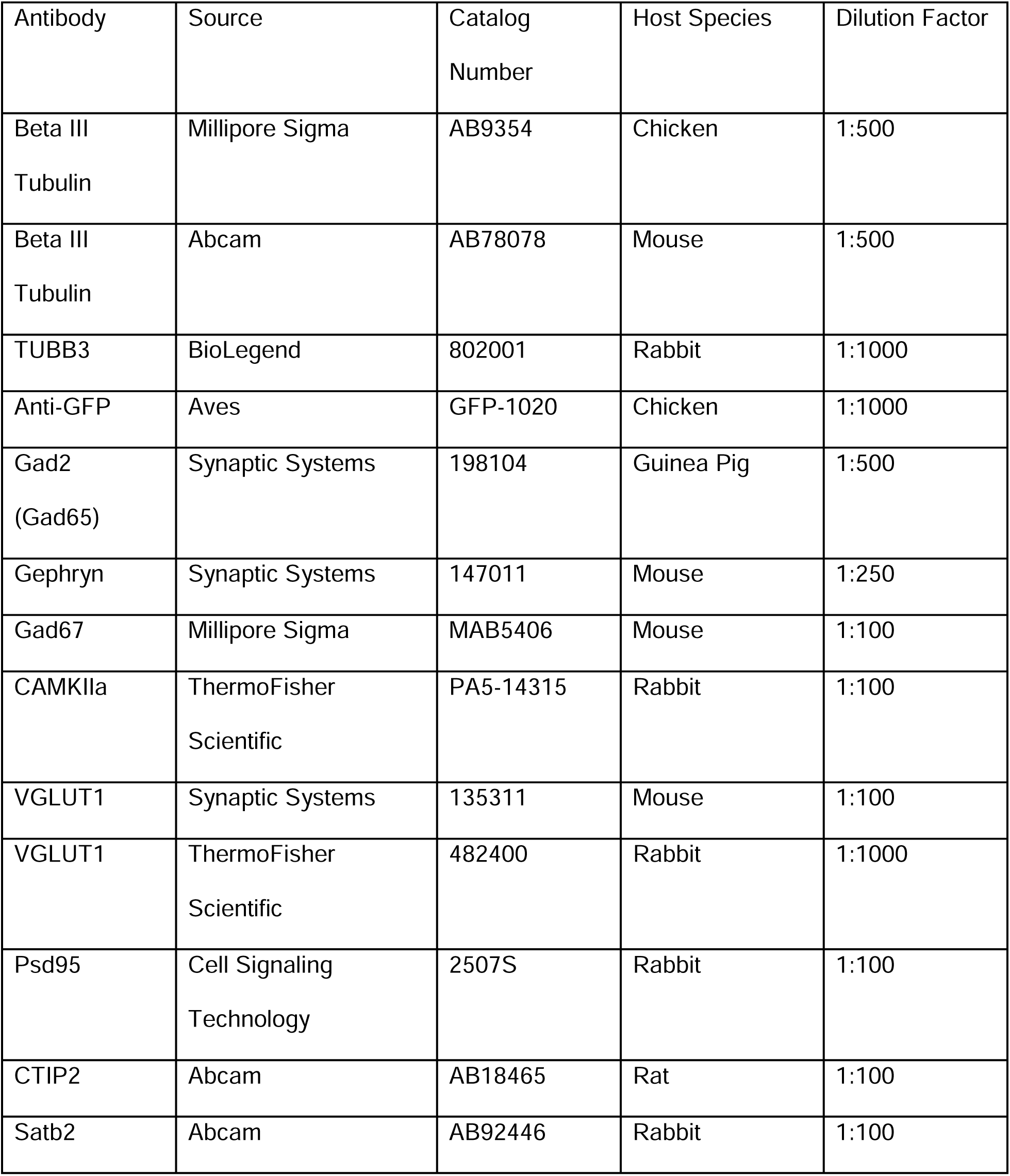
Primary Antibodies Used in Immunofluorescence Staining.

**Supplementary Table 3:** Secondary Antibodies Used in Immunofluorescence Staining.

| Antibody | Source | Catalog<br>Number | Dilution<br>Factor |
| --- | --- | --- | --- |
| Hoechst 33342 | Millipore Sigma | 34580 | 1:1000 |
| Alexafluor™ Goat anti-chicken 488 | ThermoFisher<br>Scientific | A11039 | 1:1000 |
| Alexafluor™ Goat anti-chicken 647 | ThermoFisher<br>Scientific | A21449 | 1:1000 |
| Alexafluor™ Goat anti-rabbit 488 | ThermoFisher<br>Scientific | A11034 | 1:1000 |
| Alexafluor™ Goat anti-rabbit 555 | ThermoFisher<br>Scientific | A21428 | 1:1000 |
| Alexafluor™ Goat anti-rabbit 647 | ThermoFisher<br>Scientific | A21245 | 1:1000 |
| Alexafluor™ Goat anti-rat 555 | ThermoFisher<br>Scientific | A21434 | 1:1000 |
| Alexafluor™ Goat anti-mouse 488 | ThermoFisher<br>Scientific | A11029 | 1:1000 |
| Alexafluor™ Goat anti-mouse 555 | ThermoFisher<br>Scientific | A21424 | 1:1000 |
| Alexafluor™ Goat anti-mouse 647 | ThermoFisher | A21236 | 1:1000 |
|  | Scientific |  |  |
| Alexafluor™ Goat anti-guinea pig 568 | ThermoFisher<br>Scientific | A11075 | 1:1000 |
| Alexafluor™ Donkey anti-rabbit 555 | ThermoFisher<br>Scientific | A31572 | 1:1000 |
| Alexafluor™ Donkey anti-goat 647 | ThermoFisher<br>Scientific | A21447 | 1:1000 |
| Alexafluor™ 488 conjugated<br>affiniPure donkey anti-chicken | Jackson labs | 703-545-155 | 1:250 |
| Rhodamine red-X-conjugated<br>affiniPure donkey anti-rat IgG | Jackson Labs | 712-295-153 | 1:250 |

